# Inhibition and Updating Share Common Resources: Bayesian Evidence from Signal Detection Theory and Drift Diffusion Model

**DOI:** 10.1101/2024.11.06.622265

**Authors:** Yuhong Sun, Yaohui Lin, Shangfeng Han

**Affiliations:** Department of Psychology and Center for Brain and Cognitive Sciences, School of Education, Guangzhou University, Guangzhou, China

**Keywords:** inhibition, updating, executive function, drift diffusion model, congruency sequence effect

## Abstract

Inhibition and updating are fundamental cognitive functions in humans, yet the nature of their relationship—whether shared or distinct—remains ambiguous. This study investigates the relationship between inhibition and updating within a unified task framework using a novel paradigm that integrates the N-back task with the congruent/incongruent Stroop task, creating conditions that require either updating alone or both inhibition and updating. Employing Signal Detection Theory (SDT) and the hierarchical drift diffusion model (HDDM), the results provided overall extremely strong Bayesian evidence that participants exhibited longer response times and lower accuracy in conditions requiring both inhibition and updating, compared to those requiring only updating. SDT analysis revealed a decline in discriminability, while HDDM analysis showed slower drift rates, longer non-decision times and a lower decision threshold in inhibition-demanding conditions. Even after controlling for the congruency sequence effect and current stimulus attributes, the results remained robust, showing a larger inhibition effect size compared to the traditional Stroop task. These findings suggest that inhibition consumes cognitive resources, impairing updating performance, and implying that both functions may rely on shared cognitive resources. Overall, the results elucidate the relationship between these fundamental executive functions, supporting the notion that inhibition and updating share cognitive resources.

## 1. Introduction

In the contemporary era of information abundance, knowledge acquisition is frequently impeded by irrelevant information. Consequently, it is imperative for individuals to inhibit distractions and effectively update their knowledge. Inhibition refers to the ability to overcome automatic behaviors or interference from irrelevant information to achieve a goal (Baddeley, 1996; Diamond, 2013; Miyake et al., 2000). Updating refers to the ability to receive and temporarily store information, including monitoring, adding, and deleting contents from working memory (Holder et al., 2021). While previous research has suggested that inhibition and updating are distinct components of executive function (Diamond, 2013; Espy, 2004), emerging evidence reveals correlations between performance on inhibition and updating tasks (Collette et al., 2005; Engelhardt et al., 2015; Fleming et al., 2016; Friedman et al., 2008, 2011; Ito et al., 2015; Niendam et al., 2012; Saylik et al., 2022). Thus, the relationship between inhibition and updating remains ambiguous.

The debate on whether inhibition and updating are distinct functions or share common resources persists (Jurado & Rosselli, 2007; Miyake et al., 2000). Research supporting their distinction suggested that a single-factor model combining these three executive functions (inhibition, updating, and shifting) did not fit the data as well as a three-factor model with separate components, indicating that inhibition and updating are separate processes (Miyake et al., 2000). Expanding on this, further research revealed that inhibition is more closely related to processing speed, whereas updating is predominantly linked to short-term memory capacity (Jewsbury et al., 2016). Besides, neuropsychological findings provide additional support for this dissociation. Patients with different damaged brain regions showed divergent performances in working memory and inhibition tasks, implying that these functions may be governed by different neural substrates (Godefroy et al., 1999). Saylik et al. (2022) further identified specific brain regions for inhibition and updating: the left inferior frontal gyrus for inhibition, and the bilateral middle frontal gyrus along with the left supramarginal gyrus for updating. Conversely, other evidence suggested that inhibition and updating may share common cognitive resources. For instance, Friedman et al. (2011) employed a set of independent tasks to measure inhibition and updating, revealing that inhibition and updating could be integrated into a single “Common Executive Function Factor” through latent variable analysis. More recently, large-sample studies have further supported this shared relationship (Freis et al., 2022, 2024). Consistent with this view, neuroimaging research has identified the prefrontal cortex (PFC) as a common neural substrate for inhibition and updating (Niendam et al., 2012). Despite extensive exploration, the relationship between inhibition and updating remains inconclusive.

A key reason for this ongoing debate is that researchers often measure inhibition and updating functions separately rather than concurrently. For instance, Saylik et al. (2022) used the Stroop task (Stroop, 1935) to measure inhibition and the N-back task to measure updating. Similarly, Friedman et al. (2011) used various tasks to assess inhibition and updating independently (Engelhardt et al., 2015; Fleming et al., 2016; Friedman et al., 2008, 2011; Ito et al., 2015). These traditional research approaches lack paradigms that can concurrently assess both inhibition and updating, emphasizing the need for novel experimental designs to explore their concurrent processing.

Another factor contributing to the ongoing debate is that different measurement indicators were used in inhibition and updating tasks (Von Bastian et al., 2020). Experiments often use reaction time as an indicator of inhibition and accuracy as an indicator for updating (Del Missier et al., 2010; Fleming et al., 2016; Friedman et al., 2008; Himi et al., 2019, 2021; Martins et al., 2018). A meta-analysis demonstrated that most inhibition tasks rely on either reaction time or accuracy as the sole measurement indicator, while updating tasks typically use accuracy as the primary indicator (Von Bastian et al. 2020). This discrepancy in measurement indicators reflects the distinct methods used to assess each function: inhibition tasks often use subtraction methods, while updating tasks typically use average performance (Del Missier et al., 2010; Smolker et al., 2018). Relying on a single measurement indicator poses challenges, as even within the same task, the correlation between reaction time and accuracy is low, and results vary significantly across tasks (mean *r* = .17, range = .45 to .78; Wiecki et al., 2018). This suggests that individual differences in speed-accuracy trade-offs affect the overall distribution of reaction time and accuracy. The speed-accuracy trade-off refers to the phenomenon where prioritizing one dimension (speed or accuracy) typically compromises the other (Heitz, 2014). Numerous studies have shown that reaction time is highly sensitive to speed-accuracy trade-offs (Draheim et al., 2019). Although researchers attempt to balance participants’ speed-accuracy trade-offs through instructions, participants still tend to adopt different response strategies based on their personal priorities (Draheim et al., 2021; Heitz, 2014). Additionally, individuals may adjust their speed-accuracy trade-offs to meet the demands of specific tasks (Draheim et al., 2016). Furthermore, studies have found that speed-accuracy trade-offs also occur in paradigms that solely measure accuracy (Hedge et al., 2018, 2022). Collectively, this body of evidence suggests that using a single measurement indicator is susceptible to individual speed-accuracy trade-offs, potentially distorting results. Therefore, relying on a single task with a single measurement indicator to assess specific executive functions and analyze their correlations may not accurately reflect their true relationship.

To overcome the limitation that previous studies typically measured inhibition and updating separately using different tasks, we first developed an innovative stimulus-task incompatibility experimental paradigm that enables their concurrent processing. This paradigm integrates the N-back task with the congruent/incongruent Stroop task. The N-back task requires participants to continuously judge whether the current stimulus matches the previous one (**Figure 1.A)**, thereby engaging updating (Colom et al., 2008; Hockey & Geffen, 2004; Redick & Lindsey, 2013). The congruent/incongruent Stroop task, which requires participants to classify stimuli based on whether the word’s color and semantic meaning are congruent or incongruent **(Figure 1.B)**, does not require participants to inhibit their automatic response to word meaning and therefore differs from the traditional Stroop task (Mager et al., 2007; Zurrón et al., 2009). In our study, participants were asked to judge whether the color-word congruency of the current Chinese character matched the color-word congruency of the previous one (**Figure 1.C**). This task created two critical conditions: one requiring both inhibition and updating, and the other requiring only updating. These conditions were determined by the color-word congruency of the previous trial. In both inhibition and updating required condition (when the color-word congruency of the previous trial is incongruent, e.g., ii, where the color-word congruency of both previous and current trials is incongruent; ic, where the color-word congruency of previous trial is incongruent, while current trial is congruent), inhibition was necessary to resolve the conflict between bottom-up, stimulus-driven processing (evaluating the color-word congruency of the current stimulus) and top-down, task-driven processing (determining whether the color-word congruency of the current stimulus matches the previous one). This conflict resulted in stimulus-task incompatibility, requiring participants to inhibit bottom-up processing of the stimulus attribute to successfully execute the top-down task-related judgment. In the only updating required condition (when the color-word congruency of the previous trial is congruent, e.g., cc, where the color-word congruency of both previous and current trials is congruent; ci, where the color-word congruency of previous trial is congruent, while current trial is incongruent), the bottom-up stimulus-driven processing and the top-down task-driven processing were aligned, resulting in stimulus-task compatibility. In this condition, participants could complete the updating task without inhibiting the stimulus-driven processing. Thus, our task includes conditions that require both inhibition and updating, as well as conditions that require only updating, allowing for the concurrent measurement of inhibition and updating within a unified task framework. The key assumption underlying this paradigm is that if inhibition impairs updating performance, it would indicate that inhibition and updating rely on shared cognitive resources.

**Figure 1.**
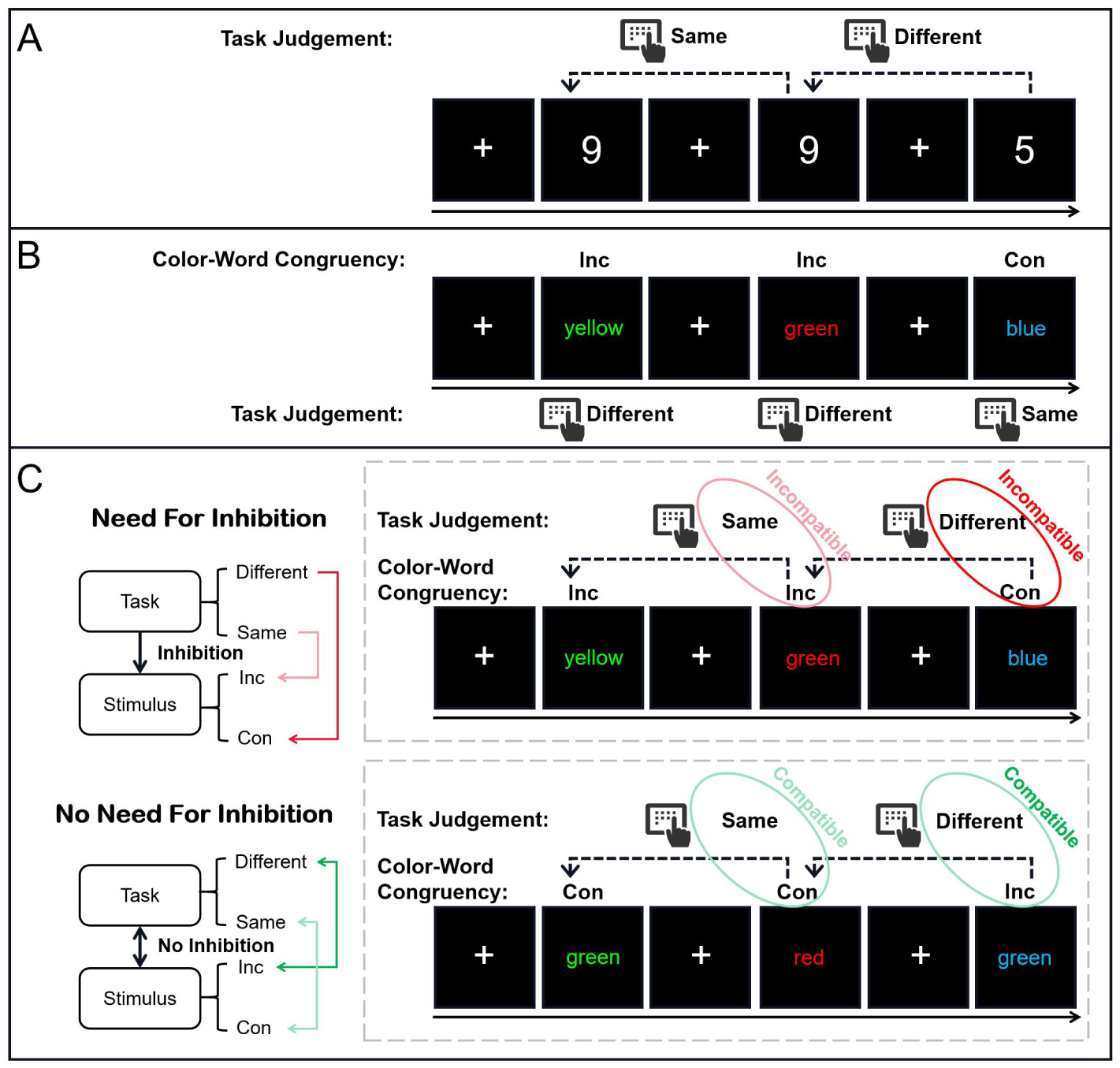
Flowchart of the Traditional N-back Task, Congruent/Incongruent Stroop Task, and Our Novel Task. (A) Traditional N-back (one-back) task. Participants are required to determine whether the current stimulus matches the previous one. (B) Traditional congruent/incongruent Stroop task. Participants are required to judge whether the current stimulus color-word congruency is congruent (same) or incongruent (different). (C) Our novel task. In each trial, a colored Chinese character is presented (For illustration, we show the English version instead of the Chinese characters). The congruency of the color and its meaning can be either congruent or incongruent. Participants are instructed to judge whether the color-word congruency of the current stimulus matches that of the previous trial. This paradigm leverages the color-word congruency of the previous stimulus to induce stimulus-task incompatibility: When the color-word congruency of the previous stimulus was incongruent (upper panel), the color-word congruency of the current stimulus conflicts with the task judgment, necessitating inhibition. When the color-word congruency of the previous stimulus was congruent (lower panel), the color-word congruency of the current stimulus aligns with the task judgment, and only needs updating.

However, our task may be influenced by inhibition-related factors. The Congruency Sequence Effect (CSE), also known as the “Gratton effect”, refers to the phenomenon where the congruency effect (typically reflected in longer response times and higher error rates on incongruent versus congruent trials) is reduced following incongruent (inhibition-required) trials compared to congruent (non-inhibition-required) trials (Duthoo et al., 2014; Egner, 2007). Previous studies have shown that CSE commonly occurs in tasks requiring inhibition (Braem et al., 2019; Frings et al., 2020). Therefore, to examine and minimize the potential impact of CSE on our results, we also considered the stimulus-task compatibility of the previous trial (i.e., whether inhibition was required in the previous trial).

Furthermore, to overcome the limitation of using different indicators for inhibition and updating tasks, this study employs Signal Detection Theory (SDT) and the hierarchical drift diffusion model (HDDM) to analyze the computational mechanisms underlying the relationship between inhibition and updating. For the N-back task, SDT is frequently utilized to evaluate updating capability. SDT analysis (Maniscalco & Lau, 2014; Stanislaw & Todorov, 1999) is a method that separately quantifies discriminability and bias through accuracy metrics. It provides a comprehensive framework for understanding how decision-makers pool information based on sensory stimuli in a bottom-up manner or how top-down information influences decisions (Green & Swets, 1966). This framework can explore whether top-down inhibition impacts updating discriminability (i.e., the discriminability index *d*’) and how the bias (i.e., decision bias *c*) to classify a stimulus as a target (or non-target) shifts under equal uncertainty conditions. A higher *d*’ shows an enhanced ability to discern if the current stimulus matches the previous one, whereas decision bias (*c*) can be interpreted as liberal (*c* < 0), neutral (*c* = 0), or cautious (*c* > 0) (Dawkins et al., 2007). If inhibition and updating share cognitive resources, simultaneous inhibition and updating should result in lower discriminability (*d*’) compared to scenarios where only updating is required, while decision bias (*c*) should remain neutral since the task does not involve probabilistic learning and the number of response keys is equally controlled, with decision bias typically linked to prior probabilities (Layher et al., 2020).

The drift diffusion model (DDM) builds upon SDT but surpasses it in certain respects. A significant advantage of DDM is its consideration of both response time and accuracy, which helps resolve issues arising from employing different measures for inhibition and updating tasks (Chwiesko et al., 2023). DDM provides a more nuanced breakdown of cognitive processes by dividing total decision time into non-decision time and decision time (Myers et al., 2022; Ratcliff, 1978; Ratcliff & McKoon, 2008), facilitating a more granular understanding of how inhibition impacts the cognitive processes underlying updating. Additionally, DDM addresses data discrepancies caused by various raw reaction time processing methods and aids in extracting parameters related to the speed-accuracy trade-off in experiments (Draheim et al., 2019; Rey-Mermet et al., 2019). According to DDM, decision-making is viewed as an evidence accumulation process, where a judgment is made once the evidence for one option meets a decision threshold (Forstmann et al., 2016). DDM breaks down reaction times on a trial-by-trial basis into underlying decision processes, represented by several psychological parameters (**Figure 2**): (1) decision threshold (*a*), which indicates the information quantity needed to make a decision and symbolizes the participants’ speed/accuracy trade-off or cautiousness. A higher threshold suggests slower yet more precise responses; (2) drift rate (*v*), which signifies the rate and potency of evidence accumulation over time; (3) non-decision time (*t*), which represents the duration of processes that are not related to decision-making (e.g., perceptual encoding of stimuli and motor responses) (Ratcliff, 1978; Ratcliff et al., 2016). A longer non-decision time can suggest the recruitment of additional attentional resources (Servant & Evans, 2020); (4) starting point (*z*), reflecting a predisposition toward one response before evidence accumulation starts (Voss et al., 2004; White & Poldrack, 2014). The hierarchical version of DDM, HDDM, has demonstrated effectiveness in estimating model parameters with fewer trials (Lerche et al., 2017; Wiecki et al., 2013).

**Figure 2.**
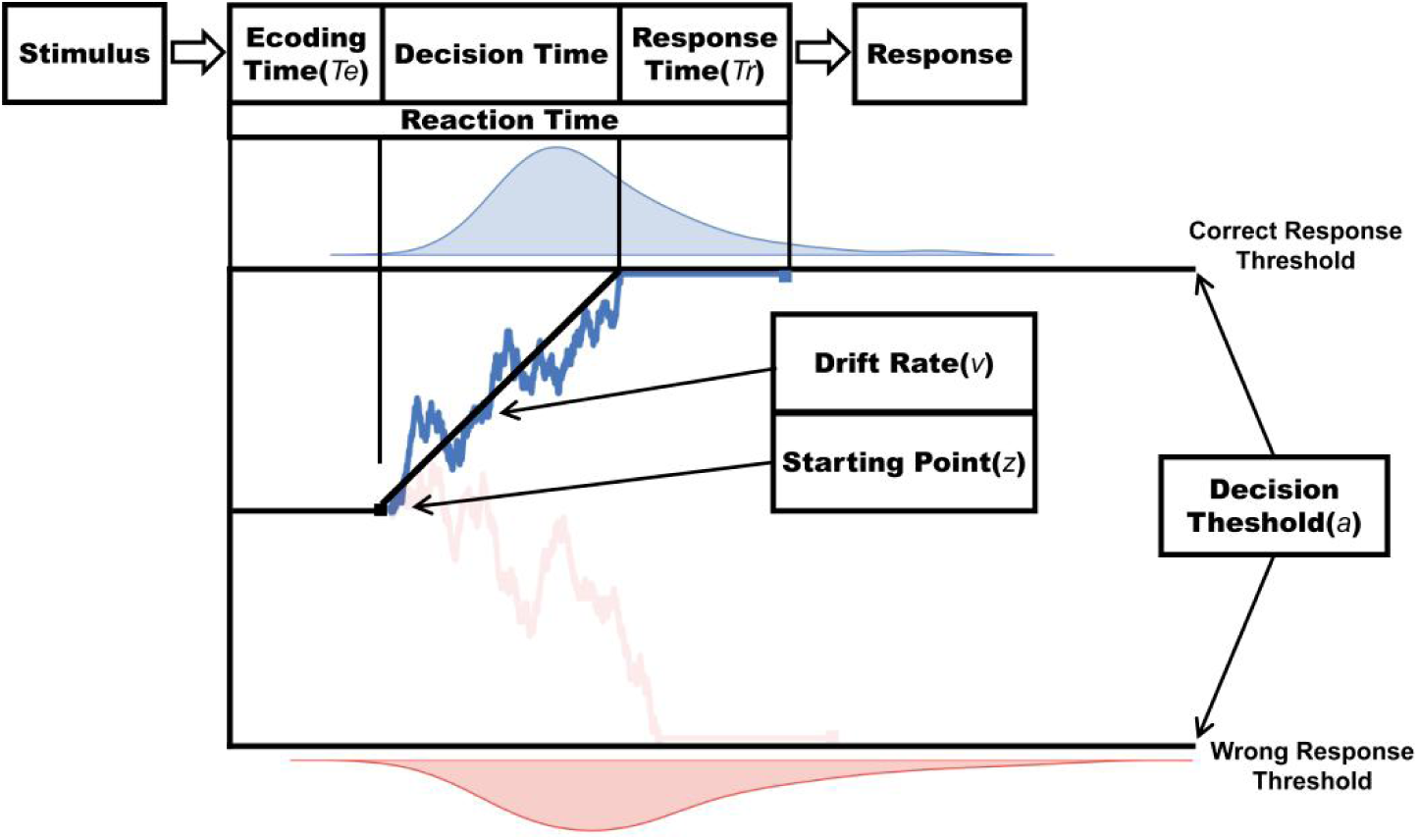
Drift Diffusion Model (DDM) Schematic.

Among the parameters in the DDM, drift rate (*v*) is a critical indicator. Prior research on the psychometric characteristics of the drift rate parameter shows that it reflects both the general cognitive ability (i.e., the speed of information accumulation necessary for completing all tasks) and cognitive processes specific to certain task conditions (Lerche et al., 2020; Schubert et al., 2016). Moreover, drift rate has exhibited good retest reliability (Lerche & Voss, 2017; Schubert et al., 2016). Based on drift rate, researchers have discovered evidence supporting the hypothesis that inhibition and updating share cognitive resources. Löffler et al. (2024) found that by evaluating drift rate across various inhibition and updating tasks, inhibition and updating were associated with a common executive function factor, implying that the two processes might share cognitive mechanisms. Thus, we hypothesize that inhibition and updating share cognitive resources. According to the dual-task paradigm (Lavie, 2005, 2010; Lavie et al., 2004; Wei et al., 2022; Wei & Zhou, 2020), if inhibition and updating do indeed share cognitive resources, performance impairments would be evident in tasks requiring both processes, leading to a lower drift rate, even when considering the CSE effect.

In conclusion, this study investigates whether inhibition and updating share cognitive resources. We propose that inhibition and updating share cognitive resources, such that when inhibition is required, updating performance is impaired even when controlling the CSE and the color-word congruency of current stimulus.

## 2. Experiment

### 2.1 Materials and Methods

#### 2.1.1 Participants

A total of 74 participants were initially recruited, but four left-handed participants were excluded from the analysis. As a result, the final sample consisted of 70 right-handed participants (39 males, 31 females), with ages ranging from 18 to 25 years (*M* = 19.90, *SD* = 1.35). All participants signed informed consent forms before the experiment. The experimental procedures were conducted in accordance with the ethical standards of the local ethics committee. Participants were rewarded with at least 15 CNY after completing the experiment.

#### 2.1.2 Materials

Stimulus materials comprised four Chinese color words (“red”, “yellow”, “blue”, and “green” in Chinese) displayed in 80-point SimSun font, with each word presented in all four colors. The RGB values were: red (255, 0, 0), green (0, 255, 0), yellow (255, 255, 0), and blue (0, 176, 240). Each image (1280 × 720 pixels) combined one of the four words with one of the four colors, yielding 16 unique stimuli. All images had a resolution of 1,280 × 720 pixels, resulting in a total of 16 (4 × 4) stimulus images. Based on the stimulus-task compatibility types of adjacent trials, eight categories were established: II (both trials involve incompatible stimulus-task), IC (previous trial involves incompatible stimulus-task, current trial involves compatible stimulus-task), CC (both trials involve incompatible stimulus-task), and CI (previous trial involves incompatible stimulus-task, current trial involves compatible stimulus-task). Additionally, the color-word congruency of the current stimulus (incongruent/congruent) was considered, resulting in 8 conditions (2 × 2 × 2 combinations). Inhibition was required for trials where there was an incompatible stimulus-task relationship (**Figure 1.C**). The experiment consisted of 198 trials, including two non-response trials at the beginning of each block. Half of the trials (98) required inhibition and updating, while the other half only required only updating. The distribution of trial types was as follows: 48 each for II, IC, and CC, and 47 for CI. Stimuli were presented in a pseudo-random order to prevent repetition priming effects (Egner, 2007), ensuring no consecutive identical stimuli were presented. Moreover, the correct response keys (“F” and “J”) were balanced across trials (Egner & Hirsch, 2005).

#### 2.1.3 Experimental Design

The experiment employed a within-subjects design with a 2 (Previous Stimulus-Task Compatibility: Incompatible vs. Compatible) × 2 (Current Stimulus-Task Compatibility: Incompatible vs. Compatible) × 2 (Color-Word Congruency of the Current Stimulus: Incongruent vs. Congruent) factorial structure. The dependent variables were reaction time and accuracy.

#### 2.1.4 Experimental Procedure

Stimuli were presented using E-Prime 2.0 (Psychology Software Tools). Each trial began with a fixation cross (“+”) displayed at the center of the screen for 1,500 ms. This was followed by a colored Chinese character, which remained on the screen for 2,000 ms or until a response was recorded. Participants were instructed to quickly and accurately judge whether the color-word congruency of the current character matched the color-word congruency of the previous trial. They were instructed to press “J” with their left index finger for “same” and “F” with their right index finger for “different”. To minimize the anticipation effect and reduce moderate variability that could influence response control (Wodka et al., 2009), a blank screen was presented for a jittered duration of 900∼1,200 ms (jittered in 100-ms increments) following stimulus presentation. The first trial of each block did not require a response (**Figure 3**).

**Figure 3.**
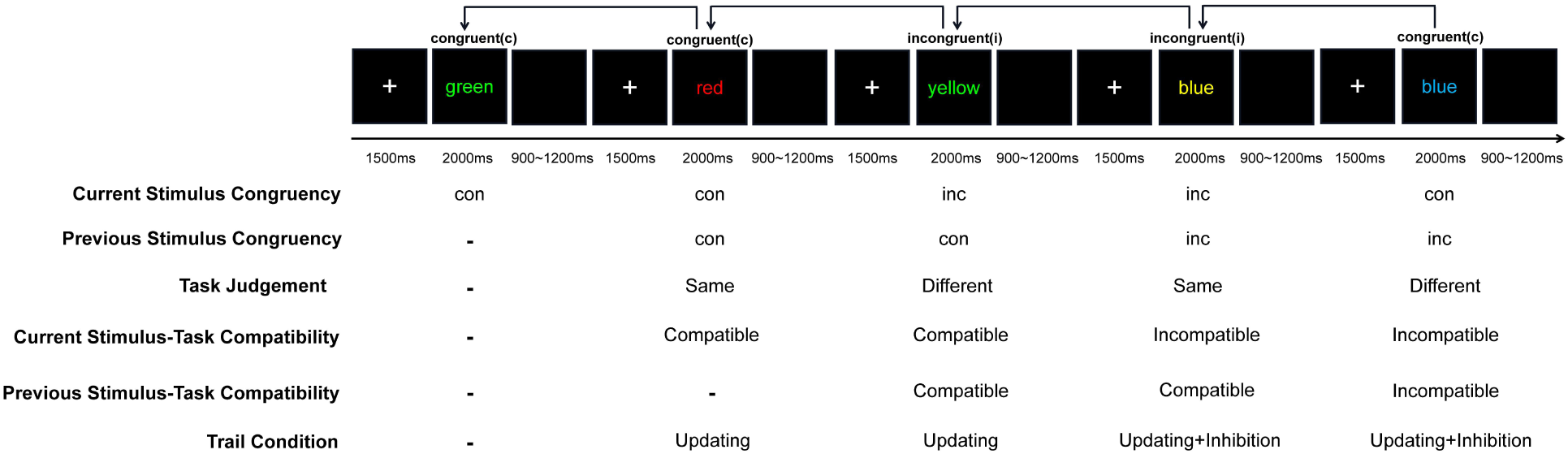
Experimental Procedure.

The temporal structure of our task ensures a fixed order of cognitive operations. Specifically, participants are required to first determine whether the current stimulus is congruent or incongruent (i.e., the congruent/incongruent Stroop judgment), and only then assess whether the type of congruency matches that of the previous stimulus (i.e., the N-back task). In the stimulus-task incompatible condition (i.e., when the color-word congruency of previous trail was incongruent; see Introduction for details), conflict arises after the congruency judgment has been completed. This sequential structure minimizes strategic variability, as the task demands inherently constrain participants to complete the congruent/incongruent Stroop judgment before engaging in the N-back comparison. Therefore, the potential for strategic task prioritization is limited by design.

Prior to the formal experiment, participants completed a practice block of 20 valid trials to familiarize themselves with the experimental procedure and key responses. Feedback on accuracy was provided at the end of the practice phase, and participants were reminded to ensure they fully understood the task before proceeding to the formal experiment. After completing 98 trials, participants were given a rest period, the duration of which was self-determined. The entire session lasted approximately 20 minutes.

#### 2.1.5 Behavioral Data Analysis

In analyzing reaction times, we excluded data from error trials and post-error trials (29.92%), as well as trials with reaction times below 200ms (0.7%), to minimize the influence of erroneous responses on subsequent trials. To examine the CSE in our task, we calculated CSE using the following formula: CSE = (CI - CC) - (II - IC).

We applied Bayesian statistics in all our analyses. Unlike *p*-values, which only indicate the likelihood of the null hypothesis being true, Bayes Factors offer a continuous measure of evidence that supports more informed decisions about hypothesis testing (Halsey, 2019; Wagenmakers et al., 2018). To examine the effects of current and previous trial types on performance, we conducted a within-subjects 2 (Previous Stimulus-Task Compatibility: Incompatible vs. Compatible) × 2 (Current Stimulus-Task Compatibility: Incompatible vs. Compatible) × 2 (Color-Word Congruency of the Current Stimulus: Incongruent vs. Congruent) Bayesian ANOVA using JASP software (V0.19.3; JASP Team, 2024) for both mean RTs and ACCs. Since no a priori information was available about the effects, we used default prior settings (*r* = .5 for fixed effects, *r* = 1 for random effects) as recommended in previous literature (Rouder et al., 2012). Bayesian analysis provides an alternative to traditional null hypothesis significance testing by offering Bayes factors. This factor quantifies the evidence for the alternative hypothesis (*H*_1_) over the null hypothesis (*H*_0_). In our study, Bayes factors were used to compare two competing models in t-tests and post-hoc tests (*BF*_10_), while inclusion Bayes factors (*BF_incl_*) evaluated the inclusion of predictors in ANOVA. Besides, we reported the standardized effect size *δ* in t-test (i.e., the population version of Cohen’s *d*, the standardized difference in mean fuse times). For parameter estimation, *δ* was assigned a Cauchy prior distribution with *r* = 1√2 (Van Doorn et al., 2021). Based on previous research on interpreting Bayes factors (Jarosz & Wiley, 2014; Jeffreys, 1998; Wetzels et al., 2011), *BF*_10_ values greater than 1 indicate stronger support for *H*_1_ , whereas values less than 1 indicate stronger support for *H*_0_ . We adopted the classification scheme for BF values outlined by Lee and Wagenmakers (Jeffreys, 1998; Lee & Wagenmakers, 2014; Speiger et al., 2025): Evidence for the *H*_1_ is considered anecdotal for a *BF*_10_ between 1 and 3, moderate for a *BF*_10_ between 3 and 10, strong for a *BF*_10_greater than 10, very strong for a *BF*_10_ greater than 30, and extreme or decisive for a *BF*_10_ greater than 100. Conversely, evidence for the *H*_0_ is considered anecdotal for a *BF*_10_ between 1 and 1/3, moderate for a *BF*_10_ between 1/3 and 1/10, strong for a *BF*_10_ smaller than 1/10, very strong for a *BF*_10_ smaller than 1/30, and extreme or decisive for a *BF*_10_ smaller than 1/100.

#### 2.1.6 SDT Analysis

Signal detection analysis was conducted, and extreme values were adjusted using the R package *psych* (Makowski, 2018). In this analysis, hits were defined as correct responses when the color-word congruency of the current and previous stimuli was the same, while misses were defined as incorrect responses under the same condition. False Alarms (FA) were defined as incorrect responses when the color-word congruency of the current stimulus differed from the previous one, and Correct Rejections (CR) were correct responses in this situation (**Figure 4**). The sensitivity index *d*’ measured updating performance, calculated as z(H) - z(F), where H and F represent the hit rate and false alarm rate, respectively. Response bias *c*, defined as the tendency to judge the current stimulus as target or non-target, was calculated as -0.5[z(H) + z(F)]. Based on our hypotheses, the signal detection analysis primarily focused on examining the interaction between current stimulus-task compatibility and previous stimulus-task compatibility to investigate the influence of inhibition on updating, while also considering the potential effect of the CSE.

**Figure 4.**
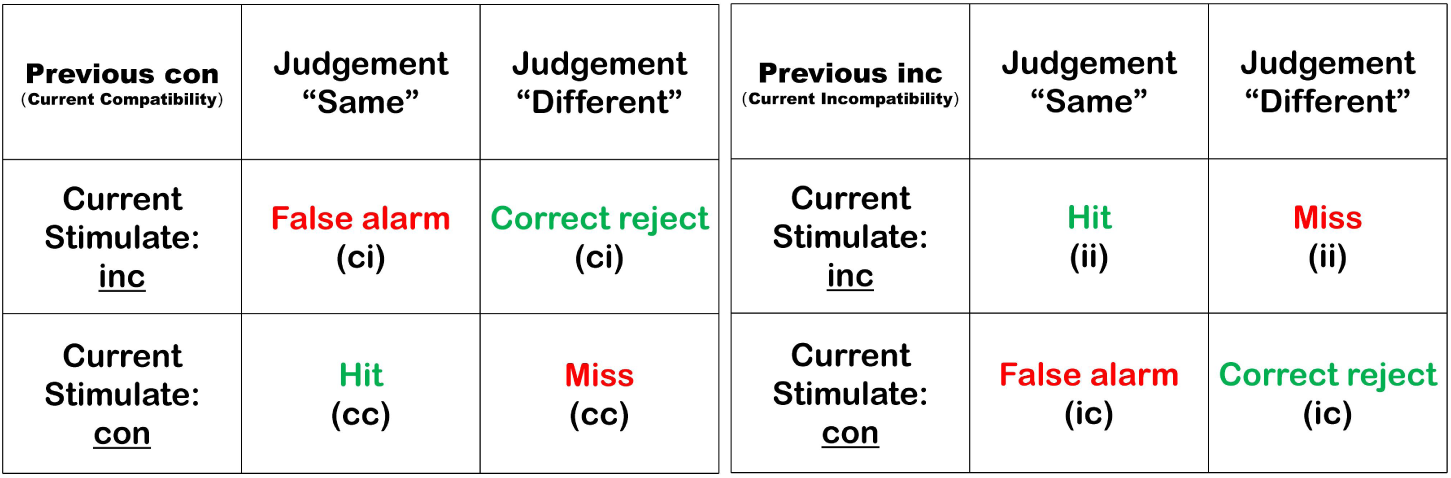
Signal Detection Theory Classification Diagram. In this diagram, green represents correct responses, and red represents incorrect responses.

#### 2.1.7 HDDM Analysis

Data modeling was conducted using the Python package HDDM (V0.9.8) within the Docker framework (Pan et al., 2025; Wiecki et al., 2013). Based on our experimental objectives and paradigm, we hypothesized that current stimulus-task compatibility would affect the drift rate, decision threshold, and non-decision time of updating. Given that color-word congruency (congruent/incongruent) is central to our task, we anticipated that color-word incongruent trials might demand more cognitive resources during the decision-making process, as participants encounter semantic conflict in these trials. Furthermore, visual properties of the current stimulus (e.g., color and brightness) were expected to influence perceptual processes during the non-decision stage (Bompas et al., 2024). Thus, we assumed that the attribute of the current stimulus would affect both drift rate and non-decision time. Regarding the potential influence of the CSE, computational modeling studies have suggested that following inhibitory trials, the activation of task-irrelevant information decreases (Koob et al., 2023; Luo & Proctor, 2022). Based on this, we predicted that the compatibility of the previous stimulus-task would impact both the drift rate and non-decision time. Since the same/different responses were balanced across trials, we set a neutral starting point (*z*) for all models. We tested four models, as outlined in Table 1.

**Table 1.**
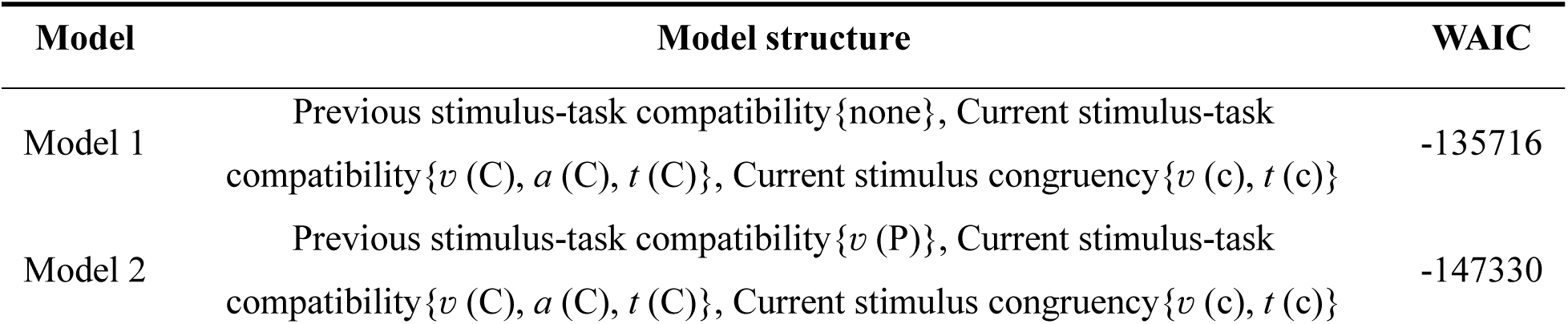
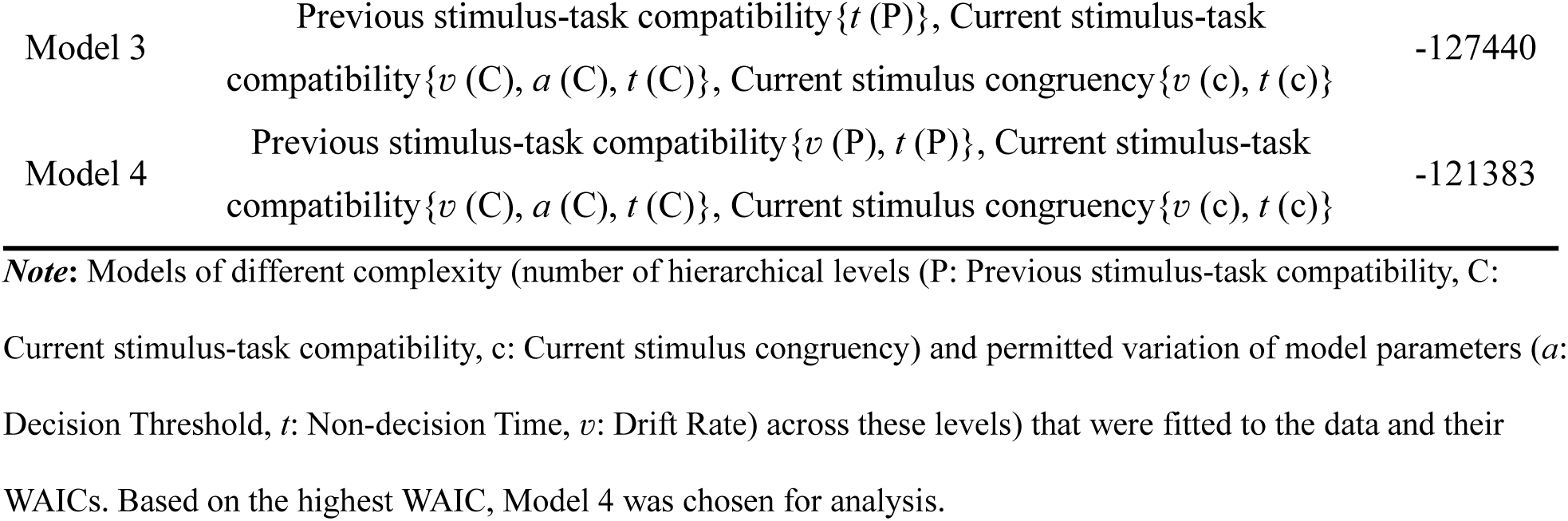
Summary of Hierarchical Drift Diffusion Models.

| Model | Model structure | WAIC |
| --- | --- | --- |
| Model 1 | Previous stimulus-task compatibility {none}, Current stimulus-task compatibility {$v$ (C), $a$ (C), $t$ (C)}, Current stimulus congruency {$v$ (c), $t$ (c)} | -135716 |
| Model 2 | Previous stimulus-task compatibility {$v$ (P)}, Current stimulus-task compatibility {$v$ (C), $a$ (C), $t$ (C)}, Current stimulus congruency {$v$ (c), $t$ (c)} | -147330 |
| Model 3 | Previous stimulus-task compatibility $\{t(P)\}$, Current stimulus-task compatibility $\{v(C), a(C), t(C)\}$, Current stimulus congruency $\{v(c), t(c)\}$ | -127440 |
| Model 4 | Previous stimulus-task compatibility $\{v(P), t(P)\}$, Current stimulus-task compatibility $\{v(C), a(C), t(C)\}$, Current stimulus congruency $\{v(c), t(c)\}$ | -121383 |

**Note:** Models of different complexity (number of hierarchical levels (P: Previous stimulus-task compatibility, C: Current stimulus-task compatibility, c: Current stimulus congruency) and permitted variation of model parameters ( $a$ : Decision Threshold, $t$ : Non-decision Time, $v$ : Drift Rate) across these levels) that were fitted to the data and their WAICs. Based on the highest WAIC, Model 4 was chosen for analysis.

Model 1 served as the baseline model, including only the effect of inhibition (current stimulus-task compatibility) and current stimulus attributes (color-word congruency of the current stimulus). Models 2, 3, and 4 were more comprehensive, incorporating the CSE (previous stimulus-task compatibility). In Model 1, we allowed drift rate (*v*), and non-decision time (*t*) to vary according to current stimulus-task compatibility and color-word congruency of the current stimulus, and decision threshold (*a*) to vary with current stimulus-task compatibility. This model aimed to explore whether inhibition and stimulus attributes influence the updating process. Model 2 extended this by allowing drift rate (*v*) to vary according to previous stimulus-task compatibility, examining the potential impact of the CSE on the decision process. Model 3, based on Model 1, allowed non-decision time (*t*) to vary with the previous stimulus-task compatibility, investigating whether the previous stimulus-task compatibility influences the non-decision phase. Model 4 expanded upon Model 1 by considering previous stimulus-task compatibility as a factor influencing both drift rate (*v*) and non-decision time (*t*).

To ensure model convergence and data validity, trials with no response or reaction times below 200ms were excluded (4.14%). For all models, parameters were estimated by drawing 10,000 samples and discarding the first 2,000. Model convergence was assessed through visual inspection of chains and calculation of the Gelman-Rubin statistic, with *R^* < 1.01 indicating good convergence (Gelman et al., 2013). For model comparison, we employed the widely applicable information criterion (WAIC; Watanabe, 2010), a fully Bayesian method that estimates out-of-sample predictive accuracy by computing the expected log pointwise posterior predictive density across the posterior distribution and applying a complexity penalty based on the effective number of parameters (Gelman et al., 2014; Vehtari et al., 2017). We calculated WAIC using the *arviz.compare* function from the ArviZ package in Python (Kumar et al., 2019), where higher values indicate better model performance on the log-score scale. All four models converged and were validated through posterior predictive checks of reaction time and accuracy (For model 4 posterior predictive checks, see **Appendix Figure 1**).

### 2.2 Results

#### 2.2.1 Behavioral Results

Regarding reaction times (**Figure 4.A, Appendix Table 1**), the Bayesian ANOVA provided extremely strong evidence for a main effect of current stimulus-task compatibility (*BF*_*incl*_ = 1.58 × 10^8^). Specifically, decisive evidence supported the reaction times for current incompatible trials (*M* = 1129.63, *SE* = 17.38) were longer than compatible trials (*M* = 1025.88, *SE* = 15.49, *BF*_10_ = 1.22 × 10^4^). Additionally, extremely strong evidence was found for the main effect of previous stimulus-task compatibility (*BF*_*incl*_ = 1120.28), where anecdotal evidence supported reaction times for previous incompatible trials (*M* = 1102.88, *SE* = 17.16) were longer than those compatible trials (*M* = 1048.92, *SE* = 14.11, *BF*_10_ = 1.60). Furthermore, there was extremely strong evidence for a main effect of color-word congruency of the current stimulus (*BF*_*incl*_ = 5.64 × 10^6^), with decisive evidence supported the reaction times for incongruent current stimuli (*M* = 1095.72, *SE* = 15.79) longer than congruent ones (*M* = 1052.48, *SE* = 15.36, *BF*_10_ = 1.04 × 10^5^), suggesting that the incongruent color-word attribute of the current stimulus leads to longer reaction times. For interaction, there was extremely strong evidence supporting two-way interaction. One between current stimulus-task compatibility and previous stimulus-task compatibility, and another between current stimulus-task compatibility and color-word congruency of the current stimulus (Both *BF*_*incl*_ ≥ 1.40 × 10^15^). In contrast, anecdotal evidence was against an interaction between previous stimulus-task compatibility and color-word congruency of the current stimulus (*BF*_*incl*_ = 0.59). Most importantly, there was extremely strong evidence supporting a three-way interaction among previous stimulus-task compatibility, current stimulus-task compatibility and color-word congruency of the current stimulus (*BF*_*incl*_ = 343.68). Post-hoc analysis revealed that the interactions between previous and current stimulus-task compatibility varied between the color-word congruency of current stimulus. In both the incongruent and congruent current stimuli, the result showed the CSE (RT CSE > 0). Pairwise post-hoc tests revealed very strong evidence that CSE was larger when the current color-word was incongruent (*M* = 503.45, *SE* = 26.27) compared to congruent (*M* = 403.45, *SE* = 27.70, *BF*_10_ = 93.40, *δ* posterior median = 0.45, 95% CI [0.20, 0.69]). Finally, to minimize the potential influence of the CSE and current stimulus attributes on the results, we analyzed only the condition where the previous trial’s stimulus-task compatibility was compatible and the color-word congruency of current stimulus was congruent, there was decisive evidence that reaction times were longer for current incompatible trials (*M* = 1255.22, *SE* = 21.04) than for current compatible trials (*M* = 847.80, *SE* = 17.61, *BF*_10_ = 3.12 × 10^29^). This result indicated that, after controlling for potential confounding factors, conditions requiring both inhibition and updating led to longer reaction times compared to those requiring only updating.

Regarding accuracy (**Figure 4.B, Appendix Table 1**), the Bayesian ANOVA provided extremely strong evidence for a main effect of current stimulus-task compatibility (*BF*_*incl*_ = 1759.87), with very strong evidence supported the accuracy for incompatible trials (*M* = 0.81, *SE* = 0.01) was lower compared to compatible trials (*M* = 0.85, *SE* = 0.08, *BF*_10_ = 36.17). In contrast, there was moderate evidence against a main effect of previous stimulus-task compatibility (*BF*_*incl*_ = 0.11) and anecdotal evidence against a main effect of color-word congruency of the current stimulus (*BF*_*incl*_ = 0.52). Moreover, there was extremely strong evidence supporting a two-way interaction between previous and current stimulus-task compatibility (*BF*_*incl*_ = 8.86 × 10^25^). Most importantly, there was very strong evidence supporting a three-way interaction among previous stimulus-task compatibility, current stimulus-task compatibility and color-word congruency of the current stimulus (*BF*_*incl*_ = 61.21). Post-hoc analysis revealed that the interactions between previous and current stimulus-task compatibility varied between the color-word congruency of current stimulus. In both the incongruent and congruent current stimuli, the result showed the CSE (ACC CSE < 0). Pairwise post-hoc tests revealed strong evidence that CSE was larger when the color-word congruency of current stimulus was incongruent (*M* = -0.36, *SE* = 0.02) compared to congruent (*M* = -0.28, *SE* = 0.02, *BF*_10_ = 10.49 , *δ* posterior median = -0.36, 95% CI [-0.60, -0.12]). Finally, to minimize the potential influence of the CSE and current stimulus attributes on the results, we analyzed only the condition where the previous trial’s stimulus-task compatibility was compatible and color-word congruency of the current stimulus was congruent, there was decisive evidence that accuracy for current incompatible trials (*M* = 0.74, *SE* = 0.02) was lower than for current compatible trials (*M* = 0.93, *SE* = 0.01, *BF*_10_ = 4.75 × 10^13^). This result indicated that, after controlling for potential confounding factors, conditions requiring both inhibition and updating led to lower accuracy compared to those requiring only updating.

**Figure 4.**
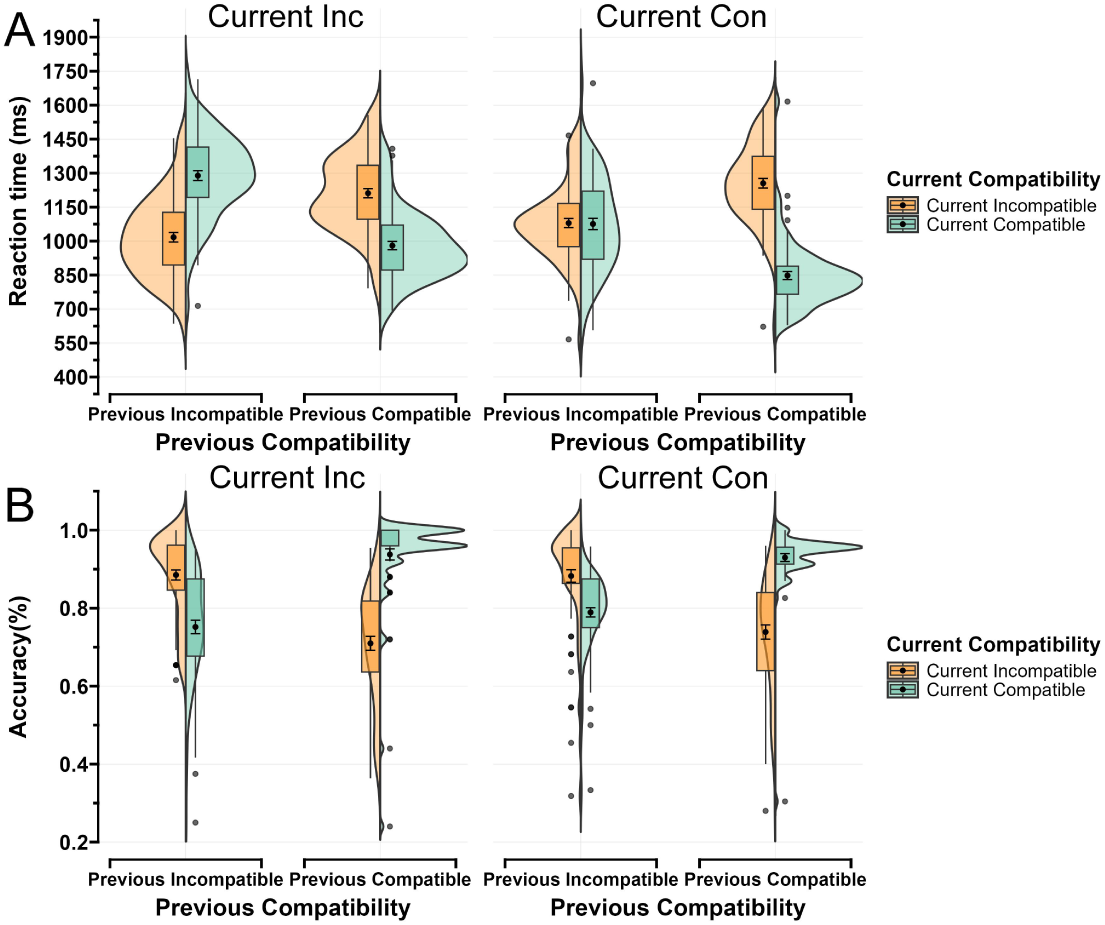
Behavioral Results. (A) Split-violin plot of reaction time. (B) Split-violin plot of accuracy. ***Note*:** In each split-violin plot, the midpoint of the box plot represents the mean of the data. The error bars around the midpoint represent the Standard Error of the Mean (SEM). The upper and lower limits of the box represent the third quartile (Q3, 75th percentile) and first quartile (Q1, 25th percentile) of the data, respectively. The whiskers extend to 1.5 times the interquartile range; data points beyond this are considered outliers. The width of the split-violin plot reflects the density of data points at different values.

#### 2.2.2 SDT Results

Regarding *d’* (**Figure 5.A, Appendix Table 2**), Bayesian ANOVA revealed strong evidence for a main effect of current stimulus-task compatibility in *d’* (*BF*_*incl*_ = 1724.87), with moderate evidence supporting that the *d’* for current incompatible trials was lower compared to compatible trials (*BF*_10_ = 3.26). In contrast, there was strong evidence against a main effect of previous stimulus-task compatibility (*BF*_*incl*_ = 0.22). Additionally, extremely strong evidence supported a two-way interaction between previous and current stimulus-task compatibility (*BF*_*incl*_ = 4.57 × 10^61^). For incompatible previous trials, pairwise post-hoc tests revealed decisive evidence that the *d’* of current incompatible trials (*M* = 2.53, *SE* = 0.12) was higher compared to current compatible trials (II vs. IC: *M* = 1.50, *SE* = 0.07, *BF*_10_ = 8.62 × 10^11^ , *δ* posterior median = 1.15, 95% CI [0.85, 1.46]). For compatible previous trials, there was decisive evidence that *d’* for current incompatible trials (*M* = 1.24, *SE* = 0.09) was lower than for current compatible trials (CI vs. CC: *M* = 3.02, *SE* = 0.09, *BF*_10_ = 1.40 × 10^26^, *δ* posterior median = -1.27, 95% CI [-1.59, -0.95]). This result indicated that, after controlling for potential confounding from CSE, conditions requiring both inhibition and updating led to lower *d’* compared to those requiring only updating.

**Figure 5.**
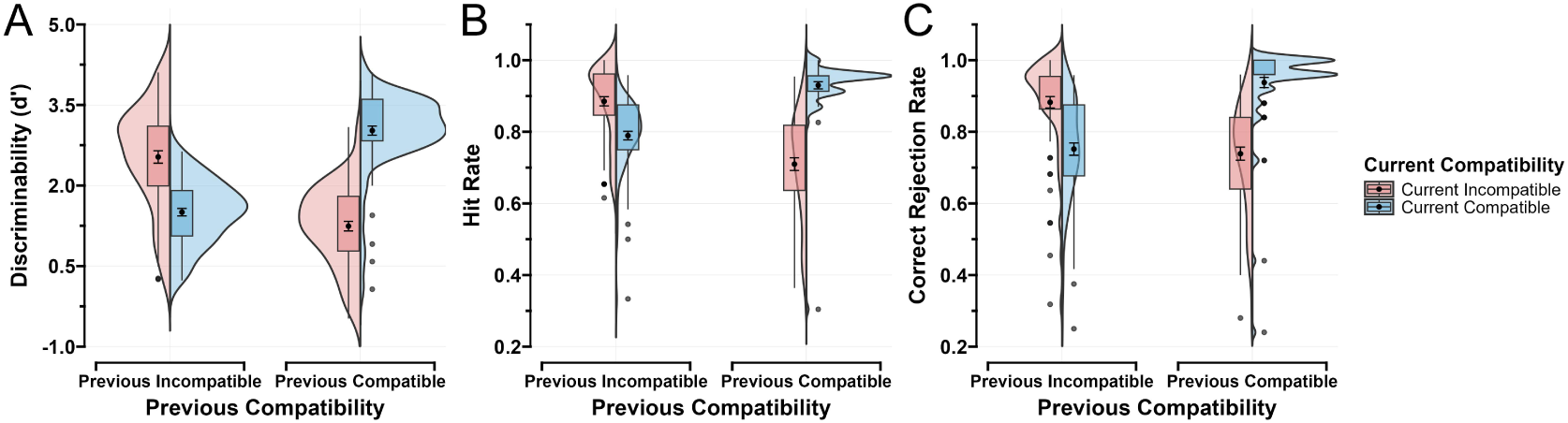
Signal Detection Analysis Results. (A) Split-violin plot of discriminability (*d*’). (B) Split-violin plot of hit rate. (C) Split-violin plot of correct rejection. ***Note*:** In each split-violin plot, the midpoint of the box plot represents the mean of the data. The error bars around the midpoint represent the Standard Error of the Mean (SEM). The upper and lower limits of the box represent the third quartile (Q3, 75th percentile) and first quartile (Q1, 25th percentile) of the data, respectively. The whiskers extend to 1.5 times the interquartile range; data points beyond this are considered outliers. The width of the split-violin plot reflects the density of data points at different values.

Regarding decision bias (*c*) (**Appendix Table 2**), Bayesian ANOVA revealed strong evidence against a main effect of current stimulus-task compatibility *c* (*BF*_*incl*_ = 0.16), but showed very strong evidence supporting a main effect of previous stimulus-task compatibility (*BF*_*incl*_ = 31.55), with very strong evidence indicating that *c* for previously incompatible trials was lower than for compatible trials (*BF*_10_ = 51.73). This suggested that experiencing conflict reduces subsequent caution in updating. Besides, there was moderate evidence against the interaction between previous and current stimulus-task compatibility (*BF*_*incl*_ = 0.55).

Regarding hit rate (**Figure 5.B, Appendix Table 2**), Bayesian ANOVA revealed very strong evidence supported a main effect of current stimulus-task compatibility (*BF*_*incl*_ = 71.17), with strong evidence supporting the hit rate for incompatible trials was lower compared to compatible trials (*BF*_10_ = 25.53). However, there was strong evidence against a main effect of current stimulus-task compatibility (*BF*_*incl*_ = 0.25). Furthermore, extremely strong evidence supported a two-way interaction between previous and current stimulus-task compatibility (*BF*_*incl*_ = 5.53 × 10^39^). For incompatible previous trials, pairwise post-hoc tests revealed decisive evidence that the hit rate of current incompatible trials was higher compared to current compatible trials (II vs. IC: *BF*_10_ = 3.04 × 10^7^ , *δ* posterior median = 0.85, 95% CI [0.58, 1.13]). For compatible previous trials, pairwise post-hoc tests revealed decisive evidence that the hit rate of current incompatible trials was lower compared to current incompatible trials (CI vs. CC: *BF*_10_ = 4.53 × 10^13^ , *δ* posterior median = -1.27, 95% CI [-1.59, -0.95]). This result indicated that, after controlling for potential confounding from CSE, conditions requiring both inhibition and updating led to a lower hit rate compared to those requiring only updating.

Regarding correct rejection (**Figure 5.C, Appendix Table 2**), Bayesian ANOVA revealed anecdotal evidence against a main effect of current stimulus-task compatibility (*BF*_*incl*_ = 0.88), and strong evidence against a main effect of previous stimulus-task compatibility (*BF*_*incl*_ = 0.29). There was extremely strong evidence supporting a two-way interaction between previous and current stimulus-task compatibility (*BF*_*incl*_ = 8.70 × 10^33^). For incompatible previous trials, pairwise post-hoc tests revealed decisive evidence that the correct rejection rate of current incompatible trials (*M* = 0.88, *SE* = 0.02) was higher compared to current incompatible trials (II vs. IC: *M* = 0.75, *SE* = 0.02, *BF*_10_ = 4.35 × 10^7^ , *δ* posterior median = 0.86, 95% CI [0.59, 1.14]). For compatible previous trials, pairwise post-hoc tests revealed decisive evidence that the correct rejection rate of current incompatible trials (*M* = 0.74, *SE* = 0.02) was lower compared to current incompatible trials (CI vs. CC: *M* = 0.94, *SE* = 0.01, *BF*_10_ = 2.15 × 10^16^ , *δ* posterior median = -1.46, 95% CI [-1.81, -1.12]). This result indicated that, after controlling for potential confounding from CSE, conditions requiring both inhibition and updating led to a lower correct rejection rate compared to those requiring only updating.

#### 2.2.3 HDDM Results

Based on the WAIC and model comparison, HDDM revealed that Model 4 provided the best fit (**Figure 6**). This suggests that the previous stimulus-task compatibility affects both drift rate (*v*) and non-decision time (*t*). Therefore, we selected Model 4 as the best-fitting model and proceeded with parameter analysis.

**Figure 6.**
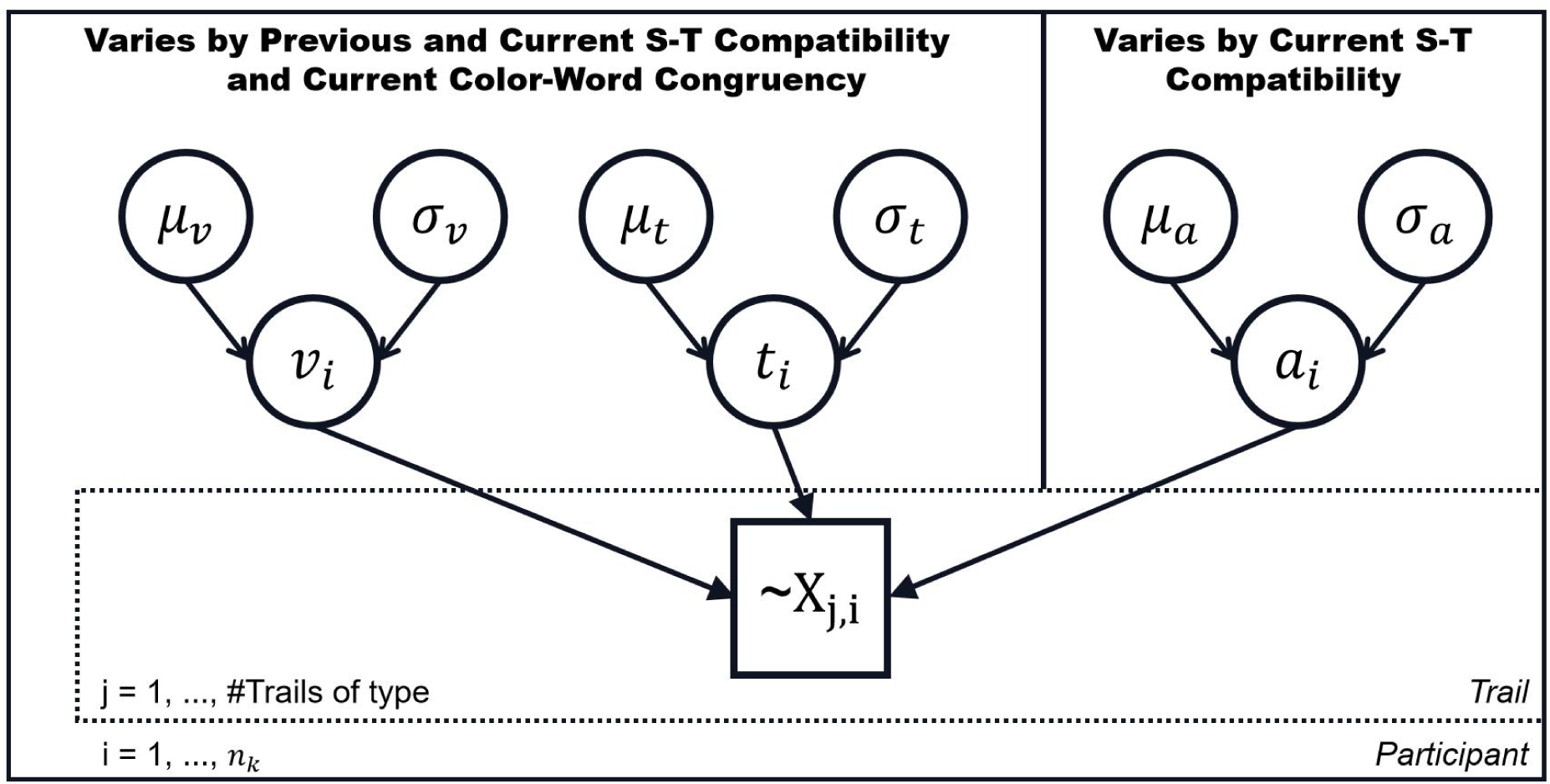
Best-fitting Hierarchical Bayesian Model Used for Estimating HDDM Parameters. The inner plate represents the participant level, while the outer plate corresponds to the group-level parameters. *v* = drift rate, *t* = non-decision time, *a* = decision threshold. Each trial is denoted as X_j,i_ . The best-fitting hierarchical model allowed drift rate, decision threshold, and non-decision time to vary by trial type. Specifically, drift rate (*v*) and non-decision time (*t*) varied based on previous and current stimulus-task compatibility and color-word congruency of the current stimulus. Meanwhile, decision threshold (*a*) was only modulated by current stimulus-task compatibility.

Drift rate (*v*) and non-decision time (*t*) were analyzed using Bayesian repeated measures ANOVA, while the decision threshold (*a*) was analyzed using a Bayesian paired samples t-test (**Figure 7, detail statistics see Appendix Table 3**).

**Figure 7.**
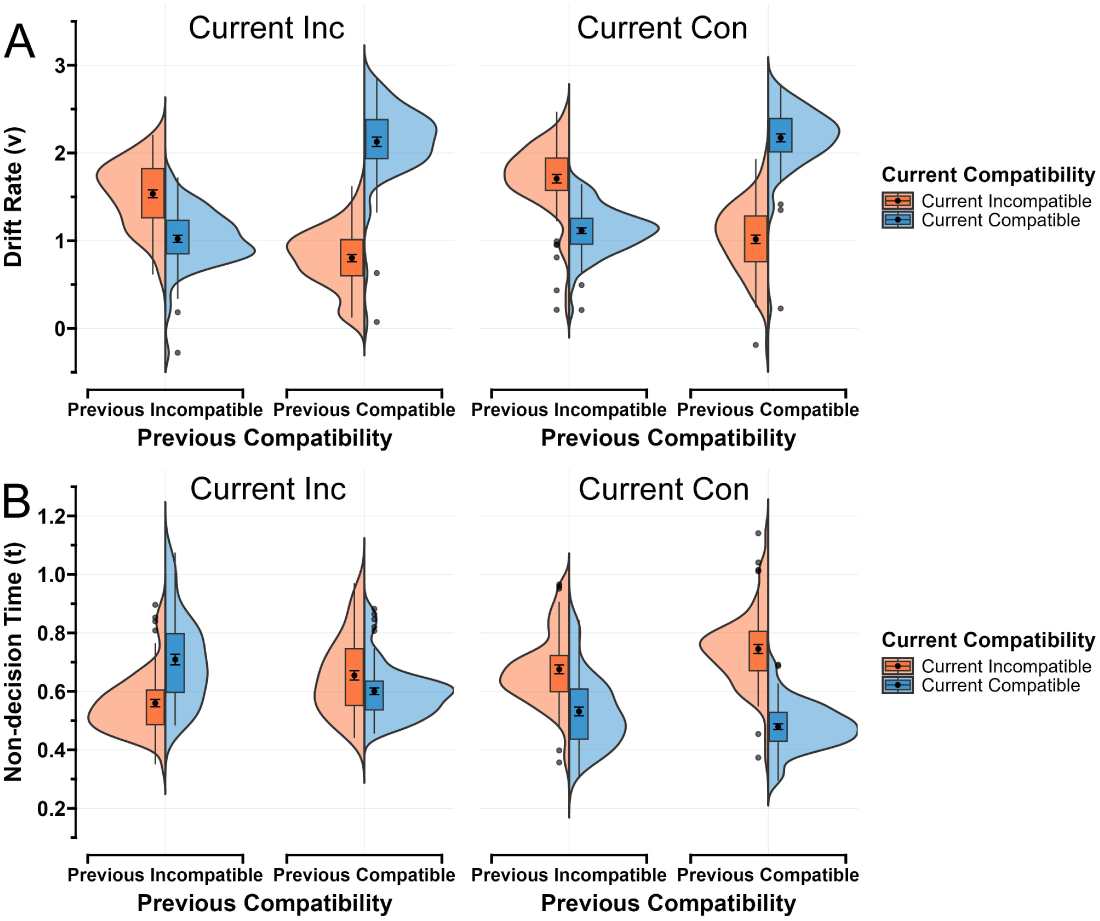
Group-level Results of Drift Diffusion Model Parameters. (A) Split-violin plot of drift rate. (B) Split-violin plot of non-decision time. ***Note*:** In each split-violin plot, the midpoint of the box plot represents the mean of the data. The error bars around the midpoint represent the Standard Error of the Mean (SEM). The upper and lower limits of the box represent the third quartile (Q3, 75th percentile) and first quartile (Q1, 25th percentile) of the data, respectively. The whiskers extend to 1.5 times the interquartile range; data points beyond this are considered outliers. The width of the split-violin plot reflects the density of data points at different values.

For drift rate (*v*), the analysis provided extremely strong evidence for the main effect of current (*BF*_*incl*_ = 2.43 × 10^16^), previous (*BF*_*incl*_ = 6.05 × 10^9^) stimulus-task compatibility and color-word congruency of the current stimulus (*BF*_*incl*_ = 1.81 × 10^5^). Specifically, decisive evidence supported the drift rates (*v*) of current incompatible trials being lower compared to compatible trials (*BF*_10_ = 4.16 × 10^5^), strong evidence supported previous incompatible trials being lower than previous compatible trials (*BF*_10_ = 9.82), and decisive evidence supported current incongruent stimuli being lower than congruent stimuli (*BF*_10_ = 7.42 × 10^5^). Furthermore, extremely strong evidence supported a two-way interaction between previous and current stimulus-task compatibility (*BF*_*incl*_ = 1.90 × 10^42^). For incompatible previous trials, pairwise post-hoc tests revealed decisive evidence that the drift rate (*v*) of current incompatible trials was higher compared to current compatible trials (II vs. IC: *BF*_10_ = 5.33 × 10^18^, *δ* posterior median = 1.64, 95% CI [1.28, 2.01]). For compatible previous trials, pairwise post-hoc tests revealed decisive evidence that the hit rate of current incompatible trials was lower compared to current incompatible trials (CI vs. CC: *BF*_10_ = 5.08 × 10^41^). This result indicated that, after controlling for potential confounding from CSE, conditions requiring both inhibition and updating led to lower drift rate compared to those requiring only updating. Besides, there was moderate evidence supporting an interaction between current stimulus-task compatibility and color-word congruency of the current stimulus (*BF*_*incl*_ = 1.93). In contrast, there was moderate evidence against an interaction between previous stimulus-task compatibility and current stimulus congruency (*BF*_*incl*_ = 0.17). Moreover, there was anecdotal evidence against a three-way interaction between previous stimulus-task compatibility, current stimulus-task compatibility and current stimulus congruency (*BF*_*incl*_ = 0.45).

For non-decision time (*t*), the analysis yielded extremely strong evidence for a main effect of current stimulus-task compatibility (*BF*_*incl*_ = 9.53 × 10^10^), strong evidence for a main effect of previous stimulus-task compatibility (*BF*_*incl*_ = 3.85), and extremely strong evidence for a main effect of color-word congruency of the current stimulus (*BF*_*incl*_ = 737.06). Specifically, decisive evidence supported the non-decision time (*t*) of current incompatible trials being shorter compared to compatible trials (*BF*_10_ = 8.50 × 10^9^), anecdotal evidence supported previous incompatible trials being longer than previous compatible trials (*BF*_10_ = 1.80), and decisive evidence supported current incongruent stimuli being longer than congruent stimuli (*BF*_10_ = 754.51). Additionally, anecdotal evidence supported an interaction between previous stimulus-task compatibility and current stimulus congruency (*BF*_*incl*_ = 2.01). There was extremely strong evidence for two way interaction between previous and current stimulus-task compatibility (*BF*_*incl*_ = 2.96 × 10^10^), current stimulus-task compatibility and current stimulus congruency (*BF*_*incl*_ = 8.15 × 10^6^). Most importantly, there was extremely strong evidence supporting a three-way interaction among previous stimulus-task compatibility, current stimulus-task compatibility, and color-word congruency of the current stimulus (*BF*_*incl*_ = 2.59 × 10^6^). Post-hoc analysis revealed that the interactions between previous and current stimulus-task compatibility varied between current stimulus congruency. In both the incongruent and congruent current stimuli, the result showed the CSE (*t* CSE > 0). Pairwise post-hoc tests revealed strong evidence that CSE was larger when the current stimulus was incongruent (*M* = 0.20, *SE* = 0.02) compared to congruent (*M* = 0.12, *SE* = 0.02, *BF*_10_ = 47.05 , *δ* posterior median =0.42, 95% CI [0.18, 0.66]). Finally, to minimize the potential influence of the CSE and current stimulus attributes on the results, we analyzed only the condition where the previous trial’s stimulus-task compatibility was compatible and the current stimulus was congruent. There was decisive evidence that non-decision time (*t*) for current incompatible trials (*M* = 0.75, *SE* = 0.02) was longer than for current compatible trials (*M* = 0.48, *SE* = 0.01, *BF*_10_ = 2.41 × 10^29^). This result indicated that, after controlling for potential confounding factors, conditions requiring both inhibition and updating led to longer non-decision time compared to those requiring only updating.

For decision threshold (*a*), the result revealed decisive evidence that the decision threshold (*a*) of current incompatible stimulus-task trials (*M* = 1.66, *SE* = 0.01) was lower than current compatible trials (*M* = 1.72, *SE* = 0.01, *BF*_10_ = 8872.37, *δ* posterior median = -0.60, 95% CI [-0.86, -0.35]).

#### 2.2.4 Exploratory Analysis: Magnitude of the stimulus-task incompatibility effects

To demonstrate the effectiveness of our manipulation of the expected inhibition constructs, we analyzed the effect sizes of the current stimulus-task incompatibility measurements and compared them with the effect sizes (Cohen’s *d*) from the previous Stroop task (Table 2).

**Table 2.** Summary of Inhibition Effect Size Presented As Mean (*SE*) [Cohen’s *d*].

| Task |  | Reaction Time | Accuracy |
| --- | --- | --- | --- |
| Our Task | Original | 103.76 (12.47) [0.99] | -0.05 (0.01) [0.56] |
|  | Control | 407.42 (18.60) [2.62] | -0.19 (0.02) [1.30] |
| Stroop | Original | 97.35 (6.01) [1.57] | -0.03 (0.01) [0.57] |
|  | Control | 104.10 (8.14) [1.25] | -0.04 (0.01) [0.49] |

When the effects of the CSE and color-word congruency of the current stimulus were not controlled (Original; comparing current incompatible vs. current compatible trials), the inhibition’s effect size for reaction time was Cohen’s *d* = 0.99, and the effect size for accuracy was Cohen’s *d* = 0.56. After controlling for the CSE and current stimulus congruency effects (Control; comparing CIc vs. CCc), the inhibition’s effect size for reaction time increased to Cohen’s *d* = 2.62, and the effect size for accuracy increased to Cohen’s *d* = 1.30. Except for the accuracy effect size when CSE and current stimulus congruency effects were not excluded (which was moderate, greater than 0.5 but less than 0.8), the effect sizes in all other cases were greater than 0.8, which is commonly considered as the threshold for large effect sizes.

We selected the Stroop task dataset^1^ from a recently published study with very strict participant selection criteria to demonstrate the traditional tasks’ effect size of inhibition (Bissett et al., 2024). For the traditional Stroop effect (Original; comparing Stroop incongruent vs. congruent trials), the inhibition’s effect size for reaction time was Cohen’s *d* = 1.57, and the effect size for accuracy was Cohen’s *d* = 0.57. After controlling for the CSE (Control; comparing Stroop ci vs. cc trials), the inhibition’s effect size for reaction time decreased to Cohen’s *d* = 1.25, and the effect size for accuracy decreased to Cohen’s *d* = 0.49. Comparing our task’s effect sizes to the traditional Stroop effect, we found that after excluding the CSE and current stimulus congruency effects, our task exhibited a larger effect size. This suggested that our task may have greater sensitivity in measuring inhibition, reflecting the effectiveness of our task in assessing inhibitory control structures. Even without excluding the CSE and current stimulus congruency effects, our task showed a comparable effect size in accuracy (Cohen’s *d* = 0.56) to the traditional task (Cohen’s *d* = 0.57), while the effect size for reaction time exceeded 0.8, indicating a large effect. Overall, our task effectively measures inhibitory control and reflects the shared relationship between inhibition and updating.

## 3. Discussion

Past research on inhibition and updating has typically examined these functions separately using distinct paradigms and different performance indicators. Such approaches often neglect the speed-accuracy trade-off, making it difficult to determine whether the relationship between inhibition and updating is shared or distinct. To address these limitations, this study investigated the relationship between inhibition and updating within a unified task framework using an innovative paradigm. By integrating signal detection theory and HDDM, we found that, regardless of the influence of the CSE, concurrent processing of inhibition impaired updating performance, suggesting that these two functions rely on shared cognitive resources. Furthermore, when comparing the inhibition effect size in our novel task with that of the traditional Stroop task, our task demonstrated a satisfactory Cohen’s *d*, further supporting its effectiveness in measuring inhibition within an updating context.

The findings support our hypothesis that inhibition and updating share cognitive resources, as evidenced by both behavioral and model-based results. Behaviorally, current incompatible stimulus-task trials resulted in increased reaction times and error rates, indicating resource competition between inhibition and updating. Signal detection analysis further provided extremely strong evidence for a decline in participants’ discriminability (*d’*) when inhibition was necessary, suggesting that cognitive resources typically allocated to updating were depleted, impairing stimulus differentiation. Computational modeling (HDDM) results further supported this view. The drift rate decreased when inhibition was needed, indicating slower evidence accumulation in the updating process. Additionally, a lower decision threshold was observed in current incompatible stimulus-task trials, suggesting that concurrent inhibition reduced the amount of evidence accumulation needed for updating, further supporting the idea that these processes share common resources. Moreover, extremely strong evidence showed longer non-decision times in current incompatible stimulus-task trials, suggesting that inhibition may have influenced the response execution stage, consuming additional cognitive resources (Bompas et al., 2024; Servant & Evans, 2020). However, potential confounds such as the CSE may have influenced our findings. To assess whether CSE was present, we analyzed the interaction between previous and current stimulus-task compatibility. For measures that exhibited only a two-way interaction between previous and current stimulus-task compatibility (*d’*, hit rate, correct rejection rate, and *v*), results consistently demonstrated the presence of CSE. Given that inhibition in one trial can affect subsequent trials and potentially influence our main findings, we excluded trials where the previous stimulus-task was incompatible and focused only on trials where the previous stimulus-task was compatible. Even under these conditions, we again found extremely strong evidence for impaired performance when inhibition was required (CI vs. CC).

Additionally, since the color-word congruency of the current stimulus could interact with CSE and influence the results, we also examined the three-way interaction between previous and current stimulus-task compatibility and current stimulus color-word congruency. The results confirmed that CSE were present regardless of whether the current stimulus was incongruent or congruent. Moreover, the CSE was stronger when the color-word congruency of the current stimulus was incongruent, suggesting that incongruent color-word congruency amplified CSE effects. These results highlight the necessity of controlling for the effects of CSE and current stimulus color-word congruency to ensure a more precise examination of the relationship between inhibition and updating. Therefore, we controlled for the condition where the previous trial’s stimulus-task compatibility was compatible and the current stimulus color-word was congruent. Even under these controlled conditions, we found extremely strong evidence for impaired performance when inhibition was required (CIc vs. CCc) in reaction time, accuracy, and non-decision time. Furthermore, after controlling the effects of the CSE and current stimulus color-word congruency, our task demonstrated an even larger effect size than the traditional Stroop task, suggesting that we successfully manipulated inhibition within the updating task. Together, these findings strongly support the notion that inhibition and updating rely on shared cognitive resources (Fleming et al., 2016; Friedman et al., 2008; Ito et al., 2015; Löffler et al., 2024). This interpretation aligns with the unity/diversity framework of executive functions (Miyake et al., 2000), which proposes that executive functions (e.g., inhibition, updating, shifting) are both separable (diversity) and intercorrelated (unity). Numerous studies using confirmatory factor analysis (CFA) have reported moderate latent correlations between inhibition and updating (Friedman et al., 2008, 2011; Friedman & Miyake, 2017; Miyake et al., 2000), suggesting that these two functions may share overlapping cognitive mechanisms. Further evidence for this unity comes from research by Löffler et al. (2024), who demonstrated that inhibition and updating are factors within a common executive function. Complementing these structural findings, causal evidence also links the two functions together. Yu et al. (2024) showed that stimulating brain areas associated with inhibition via anodal transcranial direct current stimulation (tDCS) impaired performance in updating sub-processes, indicating that disrupting inhibition undermines updating efficacy. Based on this converging evidence, we propose that shared cognitive resources may partly account for the latent correlations between inhibition and updating. Furthermore, this unity is better captured by an alternative framework in which inhibition functions as a domain-general, higher-order executive control process that is fundamentally intertwined with updating (Fleming et al., 2016; Friedman et al., 2008, 2020; Ito et al., 2015). Updating, by its nature, inherently involves inhibition, as it requires suppressing irrelevant information and removing outdated content from working memory (Zacks & Hasher, 1994).

Moreover, the lower accuracy, longer reaction times, and smaller drift rates observed when inhibition was required compared to when only updating was needed confirm that DDM can effectively capture differences between conditions (Milosavljevic et al., 2010; Ratcliff & McKoon, 2008). Additionally, the HDDM model comparison revealed that including previous stimulus-task compatibility significantly improved model fit compared to when it was excluded (Model 4 vs. Model 1). This finding suggested that the previous stimulus-task compatibility influenced both decision and non-decision processes, leading to CSE. Previous computational modeling studies have proposed that a key mechanism underlying the CSE is the reduced interference from task-irrelevant information following conflict trials (Koob et al., 2023; Luo & Proctor, 2022). Supporting this notion, our results showed that for a crucial parameter in our study—drift rate (*v*)—including previous trials’ stimulus-task compatibility led to moderate evidence against an interaction between current stimulus-task compatibility and current stimulus color-word congruency. This suggests that whether inhibition was required in the previous trial modulates the extent to which task-irrelevant information influences evidence accumulation in the current trial. Besides, we observed an interaction between the congruency of the current stimulus-task compatibility and the current stimulus color-word congruency on non-decision times, indicating that the color-word congruency of the current stimulus may affect participants’ encoding during non-decision processes.

This study provides an in-depth examination of the cognitive mechanisms through which inhibition influences updating, offering empirical support for the simultaneous processing relationship between these two executive functions. It not only introduces a new perspective on how individuals update information while resisting interference but also establishes a robust task paradigm for manipulating inhibition within an updating task. The finding that inhibition and updating shared cognitive resources highlights the importance of flexibly allocating attention and adjusting strategy selection based on task demands to optimize performance. This insight has practical implications for the development of cognitive training programs aimed at enhancing multitasking efficiency, particularly in high-stakes professions such as pilots, emergency physicians, or police officers.

Despite yielding valuable results, this study has several limitations. First, although our findings support the existence of shared cognitive resources between inhibition and updating from both behavioral and cognitive computational perspectives, previous research has found both shared and distinct cognitive resources associated with these executive functions (Engelhardt et al., 2015; Fleming et al., 2016; Friedman et al., 2008, 2011; Miyake et al., 2000; Saylik et al., 2022). The distinct mechanisms underlying inhibition and updating warrant further investigation. Second, prior studies have suggested that the N-back task may also involve proactive inhibition, specifically in suppressing irrelevant working memory information and preventing irrelevant stimuli from entering working memory (Ecker et al., 2014; Elmers et al., 2024; Kessler, 2018). However, due to the limitations of the N-back paradigm, our study could not examine how this form of proactive inhibition interacts with the inhibition manipulated in our experiment. Future research could employ paradigms such as the reference-back task to further explore these interactions (Rac-Lubashevsky & Kessler, 2016b, 2016a). This paradigm allows for the differentiation and measurement of updating sub-processes (e.g., gate opening, gate closing, substitution, and updating mode), making it possible to investigate the inhibition inherent within the updating process (Nir-Cohen et al., 2023; Yu et al., 2024). Third, our task integrated congruent/incongruent Stroop task to create an overlap in the same/different judgment attributes within N-back task. While this approach effectively introduced inhibitory demands, it also elicited semantic conflict (Mager et al., 2007; Zurrón et al., 2009), potentially confounding whether the observed effects reflect general resource depletion or modality-specific interference. Future studies could adopt non-verbal paradigms, such as spatially separated picture-word same/different judgment tasks, to enhance judgment mapping while minimizing verbal interference. Alternatively, cross-modal approaches, such as auditory-verbal tasks, may help avoid cognitive overload within a single modality and thereby clarify the source of observed performance effects.

In summary, this study employed an innovative experimental paradigm within a unified task framework, integrating signal detection theory and hierarchical drift diffusion modeling to demonstrate that inhibition and updating rely on shared cognitive resources. These shared resources primarily influence discriminability, drift rate, and non-decision time. Although the task may have been influenced by the congruency sequence effect (CSE) and the color-word congruency of the current stimulus, the results remained robust even after controlling for these potential confounds. Notably, we found extremely strong Bayesian evidence that concurrent inhibition impairs updating performance, with a larger effect size than that observed in the traditional Stroop task. These findings highlight the unity between inhibition and updating, suggesting a closely intertwined relationship between these two executive functions.

## 4. Data availability

Our raw data, analysis data, and HDDM code are accessible via the Open Science Framework: https://osf.io/t6fsu/.

## Funding

This research was supported by the MOE (Ministry of Education in China) Project of Humanities and Social Sciences (24YJC190009), Guangdong Province Philosophy and Social Sciences Planning Youth Project (GD24YXL01), Project of Guangzhou University (RC2023063), National Innovative Training Program for Chinese Undergraduate Students (202411078039).

## Contributions

Yuhong Sun: conceptualization, data curation, formal analysis, funding acquisition, investigation, methodology, project administration, software, validation, visualization, writing– original draft, writing–review and editing.

Yaohui Lin: writing–review and editing.

Shangfeng Han: conceptualization, funding acquisition, methodology, project administration, supervision, writing–review and editing.

## Ethics declarations

### Conflicts of Interest

The authors report no conflicts of interest.

## Appendix

### Appendix Tables

**Appendix Table 1.**
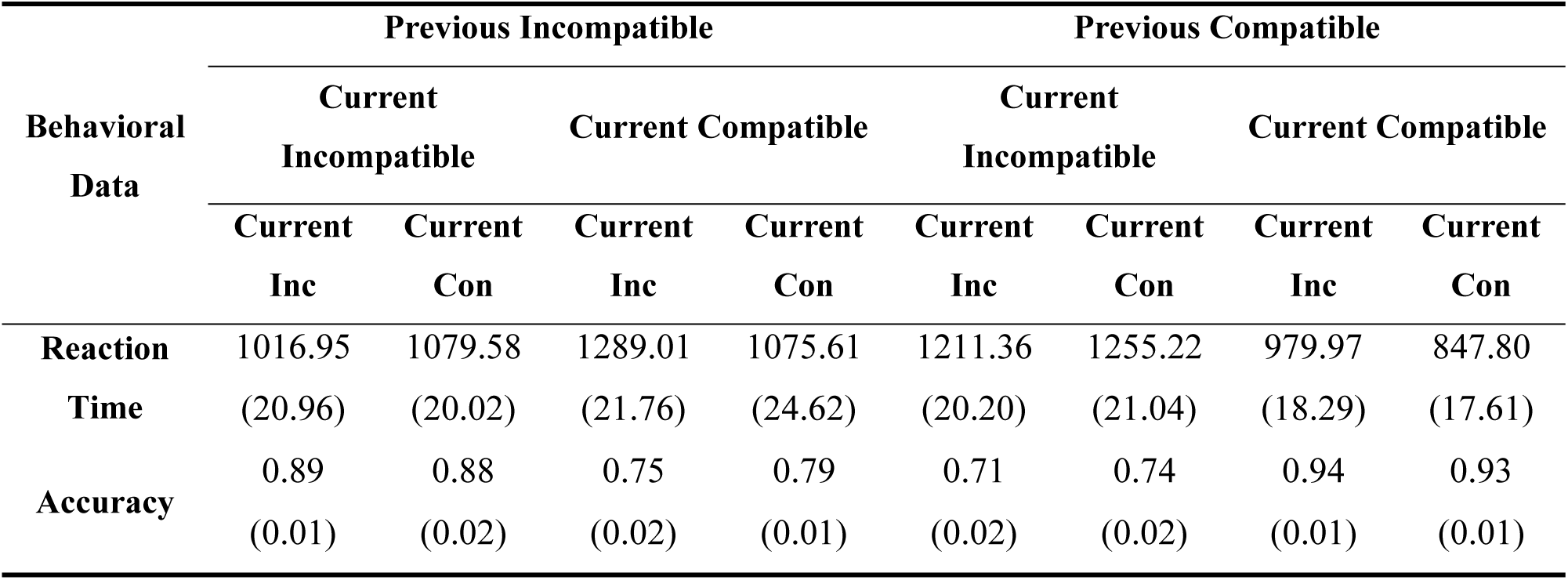
Descriptive Statistics for Behavior Data Presented As Mean (*SE*)

| Behavioral<br>Data | Previous Incompatible |  |  |  | Previous Compatible |  |  |  |
| --- | --- | --- | --- | --- | --- | --- | --- | --- |
|  | Current<br>Incompatible |  | Current Compatible |  | Current<br>Incompatible |  | Current Compatible |  |
|  | Current | Current | Current | Current | Current | Current | Current | Current |
|  | Inc | Con | Inc | Con | Inc | Con | Inc | Con |
| **Reaction** | 1016.95 | 1079.58 | 1289.01 | 1075.61 | 1211.36 | 1255.22 | 979.97 | 847.80 |
| **Time** | (20.96) | (20.02) | (21.76) | (24.62) | (20.20) | (21.04) | (18.29) | (17.61) |
| **Accuracy** | 0.89 | 0.88 | 0.75 | 0.79 | 0.71 | 0.74 | 0.94 | 0.93 |
|  | (0.01) | (0.02) | (0.02) | (0.01) | (0.02) | (0.02) | (0.01) | (0.01) |

**Appendix Table 2.** Descriptive Statistics for Signal Detection Theory (SDT) Parameters Presented As Mean (*SE*)

| SDT Parameters | Previous Incompatible |  | Previous Compatible |  |
| --- | --- | --- | --- | --- |
|  | Current | Current | Current | Current |
|  | Incompatible | Compatible | Incompatible | Compatible |
| **Hit Rate** | 0.89 (0.01) | 0.79 (0.01) | 0.71 (0.02) | 0.93 (0.01) |
| **Correct Rejection** | 0.88 (0.02) | 0.75 (0.02) | 0.74 (0.02) | 0.94 (0.01) |
| **Perceptual**<br>**Sensitivity (*d'*)** | 2.53 (0.12) | 1.50 (0.07) | 1.24 (0.09) | 3.02 (0.09) |
| **Criterion Shift (*c*)** | -0.01 (0.02) | -0.04 (0.03) | 0.05 (0.02) | 0.09 (0.03) |

**Appendix Table 3.** Descriptive Statistics for HDDM Parameters Drift Rate and Non-decision Time Presented As Mean (*SE*)

| HDDM<br>Parameters | Previous Incompatible |  |  |  | Previous Compatible |  |  |  |
| --- | --- | --- | --- | --- | --- | --- | --- | --- |
|  | Current |  | Current |  | Current |  | Current |  |
|  | Incompatible |  | Compatible |  | Incompatible |  | Compatible |  |
|  | Current | Current | Current | Current | Current | Current | Current | Current |
|  | Inc | Con | Inc | Con | Inc | Con | Inc | Con |
| **Drift Rate** | 1.53 | 1.71 | 1.02 | 1.12 | 0.80 | 1.02 | 2.13 | 2.18 |
| **(v)** | (0.05) | (0.05) | (0.04) | (0.03) | (0.04) | (0.05) | (0.05) | (0.05) |
| **Non-** | 0.56 | 0.68 | 0.71 | 0.53 | 0.80 | 0.75 | 0.60 | 0.48 |
| **decision** | (0.01) | (0.02) | (0.02) | (0.02) | (0.04) | (0.02) | (0.01) | (0.01) |
| **Time (t)** |  |  |  |  |  |  |  |  |

*Note:* Non-decision time (*t*) is measured in seconds.

### Appendix Figure

**Appendix Figure 1:**
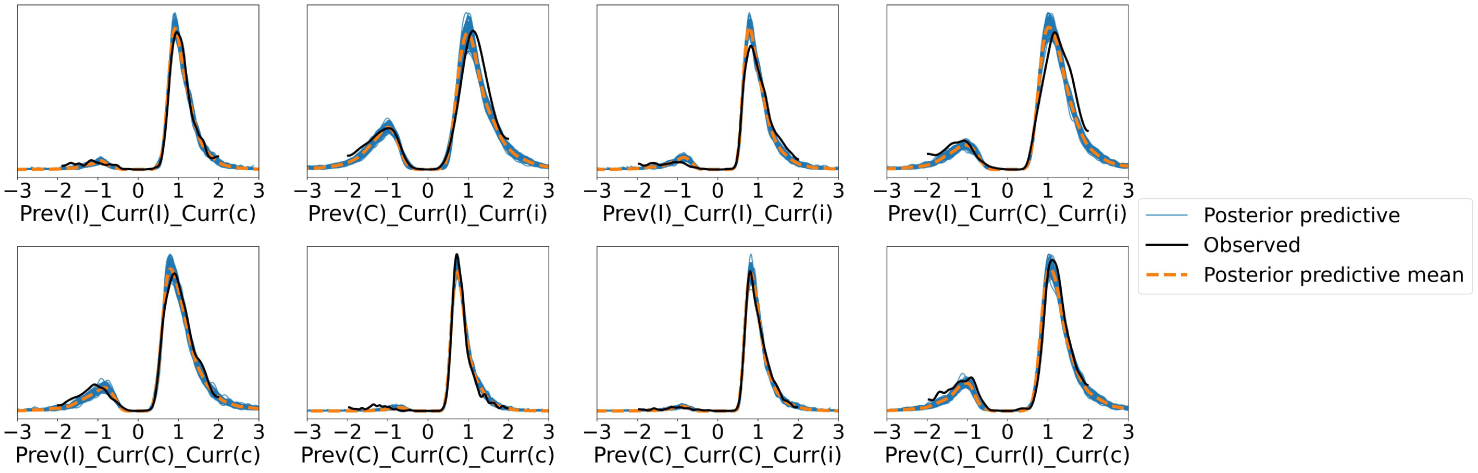
Posterior predictive check for the HDDM model 4. Blue line represent distributions of 100 randomly selected datasets generated from the posterior distribution of the model. Black line represent the distribution of observed data. Vertical axis represents probability density, and the horizontal axis represents RT (note, RT (seconds) for correct (error) responses are represented with positive (negative) values). The relative height of the negative and positive densities (divided by zero) in each panel reflects the accuracy for that condition.

## Footnotes

1 To ensure consistency in data exclusion criteria, we applied the same standards as in the current experiment. For reaction time analyses, we excluded error trials and post-error trials (9.17%), as well as trials with reaction times below 200 ms or above 2,000 ms (0.05%). Additionally, one participant with an accuracy below 50% was excluded, resulting in a final dataset of 104 participants. The compared dataset is available at our OSF link.

## Notes

https://osf.io/t6fsu/

### Competing Interest Statement

The authors have declared no competing interest.

### Summary of Updates

WAIC was used to replace DIC in the model comparison section, and some other minor modifications were made.

